# SIGHT: an immersive Virtual Reality platform for reinforced learning of optics and functional neuroanatomy of vision

**DOI:** 10.64898/2026.04.13.718109

**Authors:** Zulkifal Malik, Luca Fornia, Josef Grunig, Davide Scalisi, Federica Marchesi, Giuliano Zanchetta

**Affiliations:** Dipartimento di biotecnologie mediche e medicina traslazionale, Università di Milano, Italy -; Way s.r.l., Milano, Italy

**Keywords:** Virtual Reality, Medical Education, Optics of Vision, Functional neuroanatomy, Experiential learning

## Abstract

Virtual reality (VR) offers immersive and interactive learning environments that can improve student engagement and 3D visualization. However, its application in medical education is mostly limited to clinical settings and its potential for better understanding complex concepts, or empathy with the patients, remains underexplored. Here, we describe SIGHT (Simulated Immersive Guidance for Human Training), an immersive VR application, designed to teach core concepts in the physics and functional neuroanatomy, or neurophysiology of human vision. Its two integrated learning modules allow first-person experience of normal and pathological conditions: the optics module enables users to manipulate lenses, experience refractive errors such as myopia, presbyopia, and astigmatism and correct them through appropriate lens selection; the neurophysiology module allows learners to navigate the visual pathways from the retina to the visual cortex and to simulate lesions, experiencing the corresponding visual field deficits. User authentication and interactive evaluation steps provide analytical feedback of the experience and learning process. A pilot group of medical students reported high usability, engagement and deeper understanding of the vision-related concepts, showing how the approach of SIGHT can support experiential learning in medical education.

## Introduction

Humans learn by experiencing and interacting with their environment and deriving information from the world by using their senses (Lourenco & Casey, 2013). In virtual reality (VR) technology the sensory input derived from the real world is enhanced or replaced by sensory input created by computer simulations (Dimiter Velev, 2017). It also provides interactivity by responding to movements and actions of humans (Raptis et al., 2018). Due to its widespread potential applications to mimic reality in 3D, VR is proving to be an invaluable tool in various industries, from gaming to healthcare to education (Kamińska et al., 2019).

In recent years, VR has emerged as a transformative educational technology, by providing immersive and interactive learning experiences (Al-Ansi et al., 2023; Kamińska et al., 2019; Muñoz-Saavedra et al., 2020; Pottle, 2019; Wang, 2025). It introduces students to immersive digital experiences that traditional teaching methods cannot replicate (Phakamach et al., 2022). Therefore, it increases the engagement with complex learning materials beyond just textbooks and lectures by allowing students to experience scenarios and situations rather than imagining them (Sun et al., 2023). Studies show that VR technologies enhance students’ engagement and motivation, as well as improve their performance in academic tasks (Alizadehsalehi et al., 2021; Holly et al., 2021). Furthermore, the fun and novelty of VR learning experiences can promote interest in learning and boost learning motivation (AlGerafi et al., 2023; Liu et al., 2022). Students are more likely to invest effort when they are excited about the learning process, resulting in more favorable learning outcomes (Ou et al., 2021; Pellas et al., 2021). In medical education, thanks to its engagement and 3D visualization, VR is proving extremely useful in clinical settings, from surgery (Pelizzo et al., 2025; Rizzetto et al., 2020) to scenario based training (Davidsen et al., 2024; Mahling et al., 2023) to communication with patients (Aliwi et al., 2023). However, the potential for identification and empathy with a patient’s medical conditions is much less explored. Moreover, preclinical topics like biophysics, biochemistry, anatomy and physiology often involve complex concepts and interconnected phenomena, difficult to grasp from traditional materials. The ability of VR to support learning in these contexts remains to be proven.

The aim of the Simulated Immersive Guidance for Human Training (SIGHT) project was to develop VR learning modules for medical and medical biotechnology students about the optics and functional neuroanatomy of human vision through immersive hands-on simulations, as test cases for better understanding and reinforced learning through first-person experience of physiological and pathological conditions. Together, the two modules are expected to promote a deeper understanding of optics of vision and functional neuroanatomy of visual pathways through interactive deficit and lesion-based problem-solving, simulation, and guided experimentation. Moreover, the achievement of learning objectives was evaluated through user authentication and collection of quantitative parameters during the experience, made available to both the student and the instructor. To obtain feedback about the learning effectiveness of the VR modules, improve the quality of the teaching process and ensure a smooth user experience, we conducted a preliminary assessment with a pilot group of medical students, which provided very positive feedback.

## Methods

### Development of the pipeline

SIGHT was developed as a standalone immersive application for Meta Quest 3 using the Unity real-time 3D engine (Unity Technologies). The development pipeline combined interactive XR design, real-time physics simulation, and data analytics integration to ensure both educational effectiveness and measurable learning outcomes.

All 3D assets (laboratory environment, optical instruments, lenses, anatomical brain models, and user interface elements) were modeled and optimized using Blender and integrated into Unity for real-time rendering. The animated visual pathways and lesion simulations were implemented through custom 3D models and shader-based visual field masking to dynamically reproduce first-person perceptual deficits and the signal path from the retina towards the different regions of the brain.

Interactive mechanics (object grabbing, snapping, teleportation, hotspot activation) were implemented using Unity’s XR Interaction Toolkit, enabling controller-free hand interaction where possible and gesture-based manipulation aligned with Meta Quest 3 capabilities. The optical simulations (ray convergence/divergence, focal point detection, thin lens equation calculations) were implemented through custom C# scripts, allowing real-time computation and automated feedback based on user inputs.

The simulation of vision defects like myopia, hyperopia and astigmatism have been achieved by developing custom depth-aware post processing effects, ensuring real-time performances at the highest quality.

In-world instructional panels were developed as diegetic user interface elements anchored to spatial coordinates within the environment. These panels integrate concise text, animated diagrams, and AI-generated voice-over narration to support multimodal learning while maintaining spatial coherence and minimizing cognitive overload.

User authentication was implemented through a secure credential-based access system connected to institutional accounts. Performance metrics and user interaction data were collected via event tracking implemented in Unity and transmitted to Mixpanel (Mixpanel Inc.) for real-time analytics and dashboard visualization. The system records time-based metrics, correct vs. incorrect selections, replay frequency, and module completion data, enabling both individual and aggregated performance analysis.

The application was optimized for standalone headset performance (72–90 Hz) to ensure smooth frame rates and reduce the risk of cybersickness. Lighting was baked where possible, and asset complexity was managed through level-of-detail (LOD) optimization and occlusion culling techniques.

This technical architecture allowed the integration of immersive simulation, interactive problem-solving, and learning analytics within a single scalable XR framework.

### Feedback questionnaire

To evaluate usability and effectiveness in learning enhancement, to refine the teaching material and to identify some potential side effects of the VR experience, a pilot group of medical students at the University of Milano were recruited to test SIGHT. Participation was voluntary, and informed written consent was obtained from all participants prior to inclusion in the study. The 10 students, of both genders, were between their 2nd and 5th year and had already been exposed to the topics covered in the modules. More than half of them had some prior experience with VR devices. The students, after experiencing SIGHT, were asked to complete a questionnaire (Table 1). This was composed of 17 rating questions, on a five-point Likert scale, with responses ranging from 1 = strongly disagree to 5 = strongly agree. These were followed by 3 “Yes/No“ questions about comfort during the experience and 5 open questions to profile the users, collect their observations and identify bugs.

**Table 1.** The questionnaire used in the study.

|  |  |
| --- | --- |
| **Based on your experience, please rate your level of agreement with each of the below statements, where :1 = Strongly disagree; 2 = Disagree; 3 = Neutral; 4 = Agree; 5 = Strongly agree** |  |
| 1 | I think that I would like to use this system frequently |
| 2 | I think the system was easy to use |
| 3 | I found the system unnecessarily complex. |
| 4 | I think that I would need the support of a technical person to be able to use this system |
| 5 | I found the various functions in this system were well integrated |
| 6 | I would imagine that most people would learn to use this system very quickly |
| 7 | I found the system very cumbersome to use |
| 8 | I needed to learn a lot of things before I could get going with this system |
| 9 | I didn't have enough time to perform the activities |
| 10 | The available time was well calibrated for the activities. |
| 11 | The experience was too long |
| 12 | Virtual reality made learning more engaging than a classroom lecture |
| 13 | The VR experience helped me better understand the concepts covered |
| 14 | I found the visual representation of the phenomena (eye, light rays, neural pathways) clear and intuitive. |
| 15 | I didn't learn anything new compared to traditional lectures |
| 16 | I believe VR will help me retain the concepts I've learned longer |
| 17 | I would prefer more university courses to include VR activities |
| **Yes/No Questions** |  |
| 18 | Did you experience any headache? |
| 19 | Did you experience any dizziness? |
| 20 | Did you experience any eye strain? |

Table 1. The questionnaire used in the study
| Open Questions |  |
| --- | --- |
| 21 | Which aspect of the VR experience did you find most useful or interesting? |
| 22 | What difficulties did you encounter, if any? |
| 23 | Do you have any suggestions for improving the experience or the learning content? |
| 24 | Which academic year of study are you in? |
| 25 | Do you have previous experience with VR, for educational purposes, or any other purpose? |

After collecting the questionnaires, the full data (Suppl. Fig. 1) were analyzed to determine the respondents’ opinions, clustering questions for usability (questions 1-8), appropriateness (9-11), engagement (12) and learning effectiveness (13-17). The score for each category was obtained as the average of the corresponding scores (using an inverse scale for the negative questions).

## Results

### General design principles and distinctive features

We have designed an immersive VR platform for medical and biotechnology students, for enhanced learning through interactive modules and first-person experience of concepts and phenomena related to human physiology and relevant defects. The first two integrated modules focus on vision (Fig. 1). The first module addresses optics, enabling the user to explore lens types (converging and diverging) and experience common refractive errors, including myopia, presbyopia and astigmatism, and their correction. The second module focuses on functional neuroanatomy, or neurophysiology and provides a guided, 3D visualization of the visual pathway from the retina to the visual cortex. The user can interactively apply lesions at different points along the pathway, experiencing the resulting visual field defects in real time. More details about the modules will be provided in the next sections.

**Figure 1.**
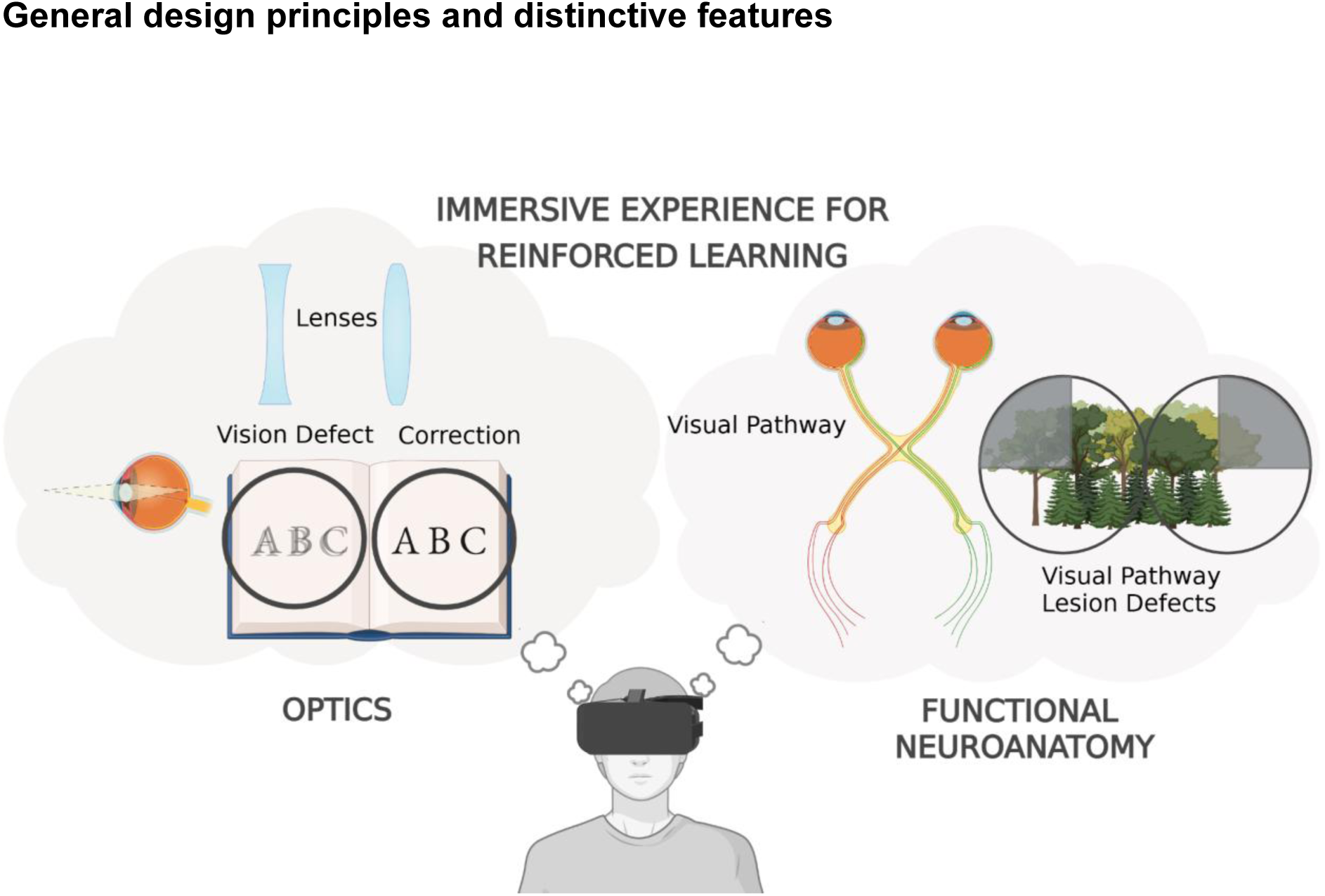
General structure of SIGHT and content of the two integrated modules.

The overall user flow follows a structured sequence consisting of a tutorial area, authentication, and module selection with onboarding. Several core components were conceived to support user interaction and learning within the VR environment. The application modules are delivered within a modern laboratory to provide a familiar and intuitive learning space. Interactive tasks are organized across multiple workstations. Each workstation contains a clearly marked hotspot that enables the user to activate tools and manipulate objects required for task completion. The user can move freely within the environment; however, approaching a hotspot triggers automatic positioning at the corresponding workstation to facilitate interaction.

#### Tutorial Area

Each time the application is launched (Fig. 2A), users are presented with a brief, interactive tutorial designed to introduce core VR mechanics, where the users familiarize themselves with the controls and navigation within the VR environment; the tutorial can be skipped at any time. The tutorial includes hand gestures to grab by pinching, touch by poking, aim and pinch to teleport. It is based on a simple activity, inserting test tubes in a rack by pinching and grabbing, to make users more confident with the gestures (Fig. 2B, C).

**Figure 2.**
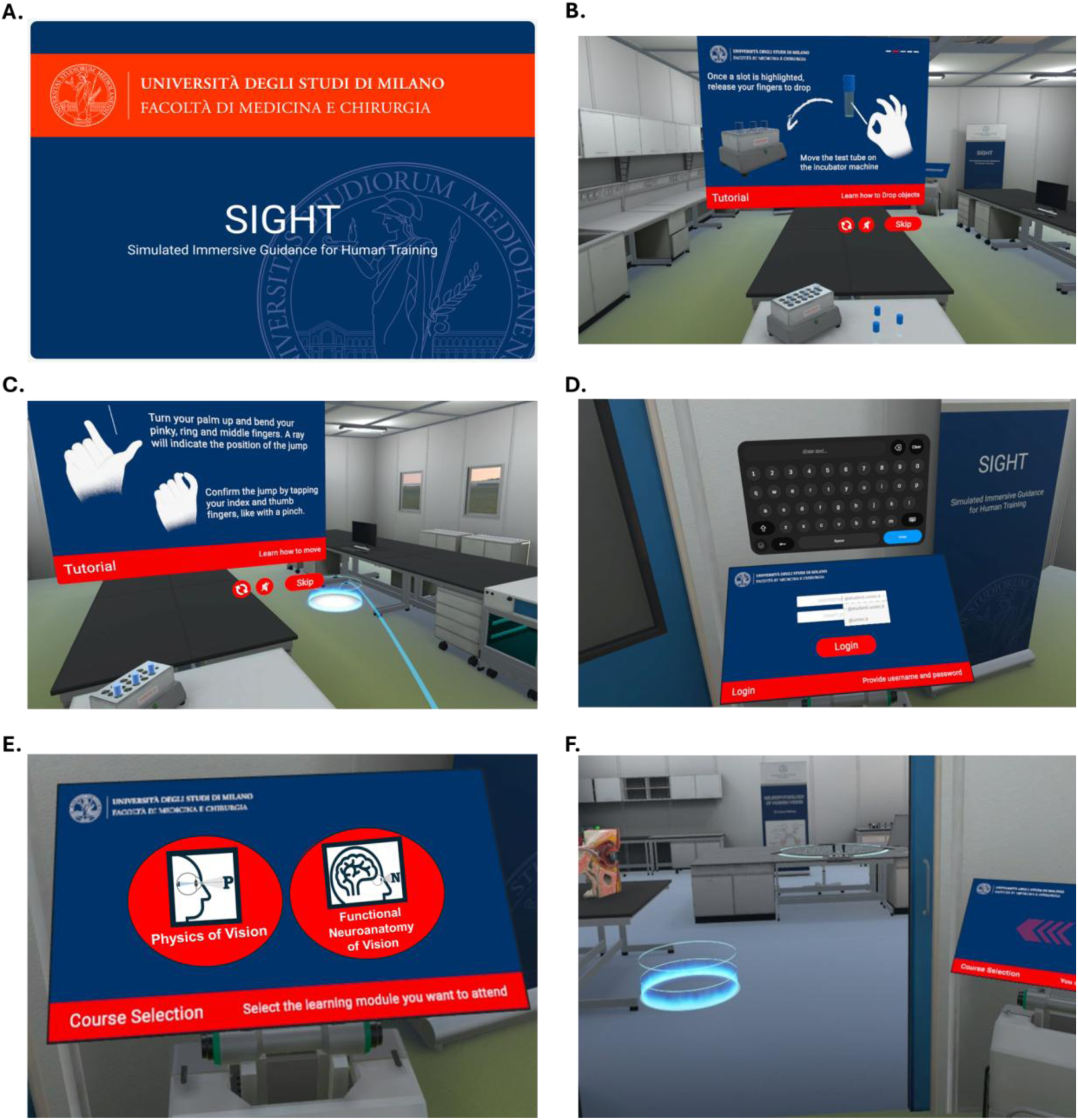
Preliminary activities. A. SIGHT screen welcomes the user at the beginning of the experience. B. A tutorial shows basic hand gestures to interact, grab and move virtual objects. C. The primary locomotion system, teleportation through hand gestures, is explained. D. Authentication with University credentials. E. Users can select the module. F. The entrance of the laboratory where the activities are performed.

#### Authentication, module selection and onboarding

An authentication interface was implemented to allow users to log in using their University of Milano credentials, enabling user-specific access and data tracking. Log in is performed on a diegetic user interface in proximity to the entrance door of the laboratory (Fig. 2D). Users are then presented with a module selection interface that allows them to choose between the available learning modules (Fig. 2E). Once a module is selected, the sliding door opens and the user is guided into the laboratory (Fig. 2F). Educational information and instructions are delivered through in-world pop-ups and floating panels that appear contextually near objects or tools the user is interacting with. These panels include concise text, icons, AI-generated voice-over, and, where appropriate, animations or short videos (e.g., anatomical illustrations or optical diagrams). Panels are positioned within an optimal viewing range and automatically disappear when no longer relevant to minimize visual clutter.

#### Data collection and feedback

User-specific performance and interaction data are collected throughout the experience. Metrics such as time spent on tasks, accuracy, and total session duration are recorded and transmitted to the analytics platform for aggregation and analysis. Upon completion of the experience, users are provided with feedback summarizing their activity and performance.

### Physics of human vision module

The Physics of Human Vision module consists of three interactive experiences designed to teach lens types, refractive errors (myopia, presbyopia, astigmatism), and their correction (Fig. 3).

**Figure 3.**
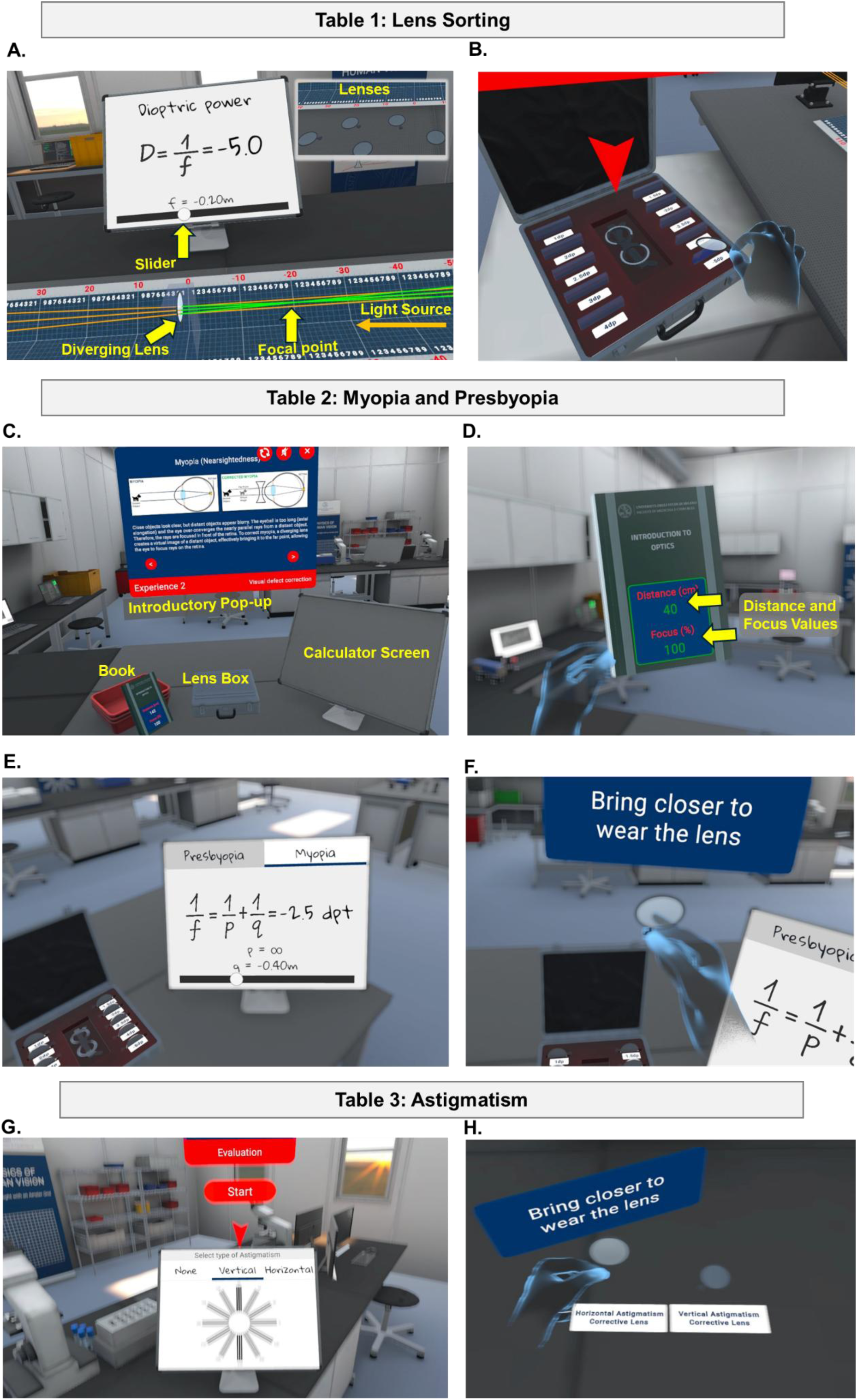
Optics module. A. When a diverging lens is placed along parallel light rays, a virtual focal point is determined by the intersection of virtual (green) rays. The corresponding dioptric power is calculated with the aid of the interactive calculator. B. The lenses are inserted into their corresponding slots in a briefcase. C. The setup for defect assessment and correction, with introductory information about myopia, hyperopia and presbyopia. D. A randomly applied visual defect can be quantified by measuring the threshold distance with the display on the book. bringing a book closer or farther. E. The whiteboard calculator, based on the thin lens equation, provides the estimate for the required lens. F. The lens is brought to the eyes and feedback is provided about the selection. G. Vertical and horizontal astigmatism can be applied and experienced. H. In the evaluation phase a random vertical or horizontal astigmatism can be corrected by bringing the appropriate lens close to eyes.

In the first table, students are introduced to converging and diverging lenses and tasked with classifying a group of lenses based on their dioptric power. When the users insert a lens in the path of parallel light rays, these converge for convex lenses and diverge for concave lenses, with a virtual, negative focal point (Fig. 3A). The focal length *f* is measured with a ruler and entered into a calculator to obtain the dioptric power as *D =* 1*/f.* A total of 10 lenses are included in this task, with dioptric powers from +1 to +4 and from −1.5 to −5, respectively. The students must place all the lenses into their correct slots in a briefcase (Fig. 3B) and upon completion they are invited to proceed to the next table. Throughout the experience, students are guided with the help of voice-overs and panels, appearing after mistakes or in case of inactivity, to clarify concepts or suggest how to proceed.

The second table is devoted to the basic principles of myopia and presbyopia, virtually experienced by the users, and their correction using appropriate lenses. Introductory information about the visual defects and their correction is presented through contextual pop-ups (Fig. 3C). The workstation includes a set of lenses, an interactive whiteboard and a book that provides real-time feedback on viewing distance and on image crispness (Fig. 3D). A randomly generated visual defect is then applied to the student, from a range of values for far point (20-67 cm) in myopia and near point (25-100 cm) in presbyopia. Users can identify and quantify the defect by picking up the book and adjusting its distance from the eyes. The simulation reproduces the characteristic visual impairments: distant objects appear blurred in myopia, whereas near objects appear blurred in presbyopia. Through the dynamic indicators on the book cover, the near (far) point can be determined. Users must then select the appropriate thin lens equation on the whiteboard, inserting the measured distance and calculating the required dioptric power for correction (Fig. 3E). The proper lens is selected from the available set and applied by bringing it toward the eyes (Fig. 3F). Correct selection restores normal vision, whereas incorrect choices trigger immediate visual feedback, prompting further attempts. If repeated errors occur, additional guidance is provided through a “help” function that includes solved examples demonstrating the application of the thin lens equation. Upon successful correction, users may proceed to the next task or repeat the activity with a different defect type or severity.

The third table introduces the main types of astigmatism through schemes and a video in a contextual panel. Users then virtually experience this condition while looking at a grid pattern (Fig. 3G) or the surrounding environment. In the final evaluation task, a randomly generated astigmatism is applied and must be identified and corrected by wearing the appropriate cylindrical lens from the available options (Fig. 3H).

### Functional neuroanatomy of human vision module

This module introduces students to the visual pathways from retinal input to the primary visual cortex passing through the lateral geniculate nucleus in the thalamus and allows to test how lesions at different levels of these pathways lead to specific patterns of visual impairment.

In the first part, after a pop-up informs about the activity to be carried out, the user is presented with a semi-transparent 3D rendering of the brain, incorporating a detailed reconstruction of the visual pathways, from eyes to the calcarine fissure in the bilateral occipital lobe, where area 17, the primary visual area, is located (Fig. 4A). The model is displayed on an interactive turntable and can be zoomed and freely rotated along multiple axes via two joysticks. Ten hotspots highlight specific critical nodes within the pathway, which the student can select and learn about (Fig. 4B and Suppl. Table 1). Afterwards, it is possible to select and apply simulated focal lesions in various anatomical locations and experience the corresponding visual field deficit for a few seconds (Fig. 4C,D and Suppl. Table 2). The user can try all defects and then proceed to the second part of the module, where the acquired knowledge is assessed (Fig. 4E). A random visual field deficit is directly experienced and its visualization is shown on the whiteboard nearby. The student is asked to identify the precise anatomical location of the lesion, responsible for the experienced impairment (Fig. 4F). If the wrong area is selected, the field deficits are compared in the whiteboard and the user is invited to try again (Fig. 4G) while, if the correct lesion is selected, the student can repeat the experience with a different defect. After three successful damage selections, the evaluation metrics are shown to the user and sent to the instructor (Fig. 4H).

**Figure 4.**
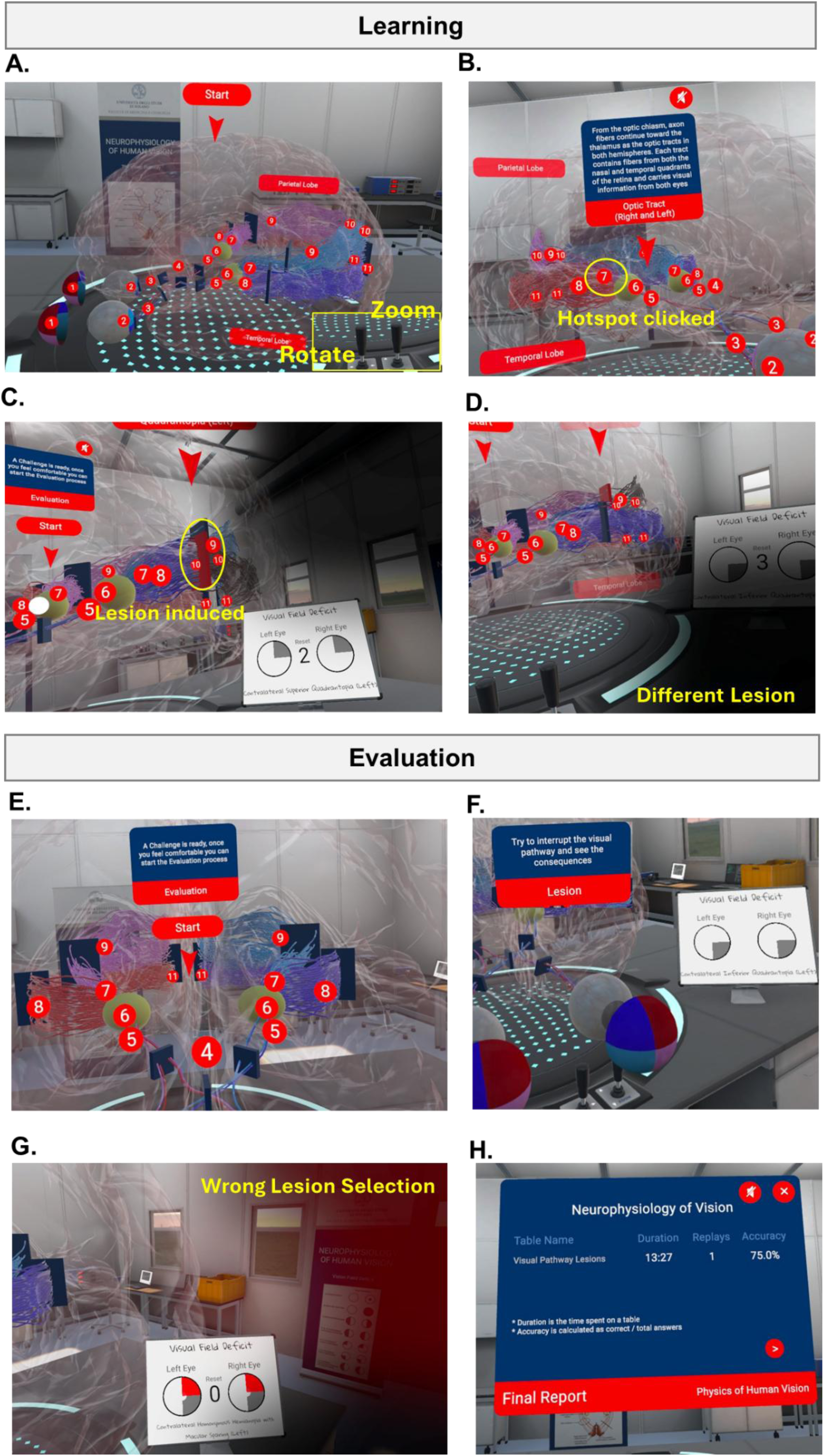
Functional neuroanatomy module. A. 3D view of the brain, with the main nodes of the visual pathways. B. By clicking on a hotspot, a pop-up appears with a description of the node. C-D. Lesions can be applied by clicking on the corresponding position along the visual pathway, allowing the user to experience their consequence on the visual field. E. Start of the evaluation activity. F. A defect is shown on the screen and the correct lesion spot must be identified. G. When a wrong lesion is selected, an error message appears and visual field impairments are compared. H. At the end of the module, students are shown quantitative indicators of their activities.

### Preliminary student assessment

To get a preliminary evaluation of system usability and user feedback, SIGHT was proposed to a pilot group of medical students of the University of Milano on the Meta Quest 3 head-mounted device. The students tested both modules and were asked to complete a questionnaire (Table 1) at the end of the experience. The full answers (reported in Suppl. Fig. 1) were clustered to extract overall indicators about the VR experience and the perceived educational value. The questionnaire, on a 1-5 scale, revealed an average score of 4.2 assigned to the system usability, 4.5 for the appropriateness of the allocated time and 4.8 both for the engagement and the learning effectiveness (Fig. 5). Overall, students witnessed a very positive experience, which they would like to extend to other topics and courses. As regards issues related to the VR experience, some of them reported dizziness (40%), eyestrain (20%) or headache (10%). The comments in the open questions suggested minor changes to optimize the user experience, which were implemented in the following version.

**Figure 5.**
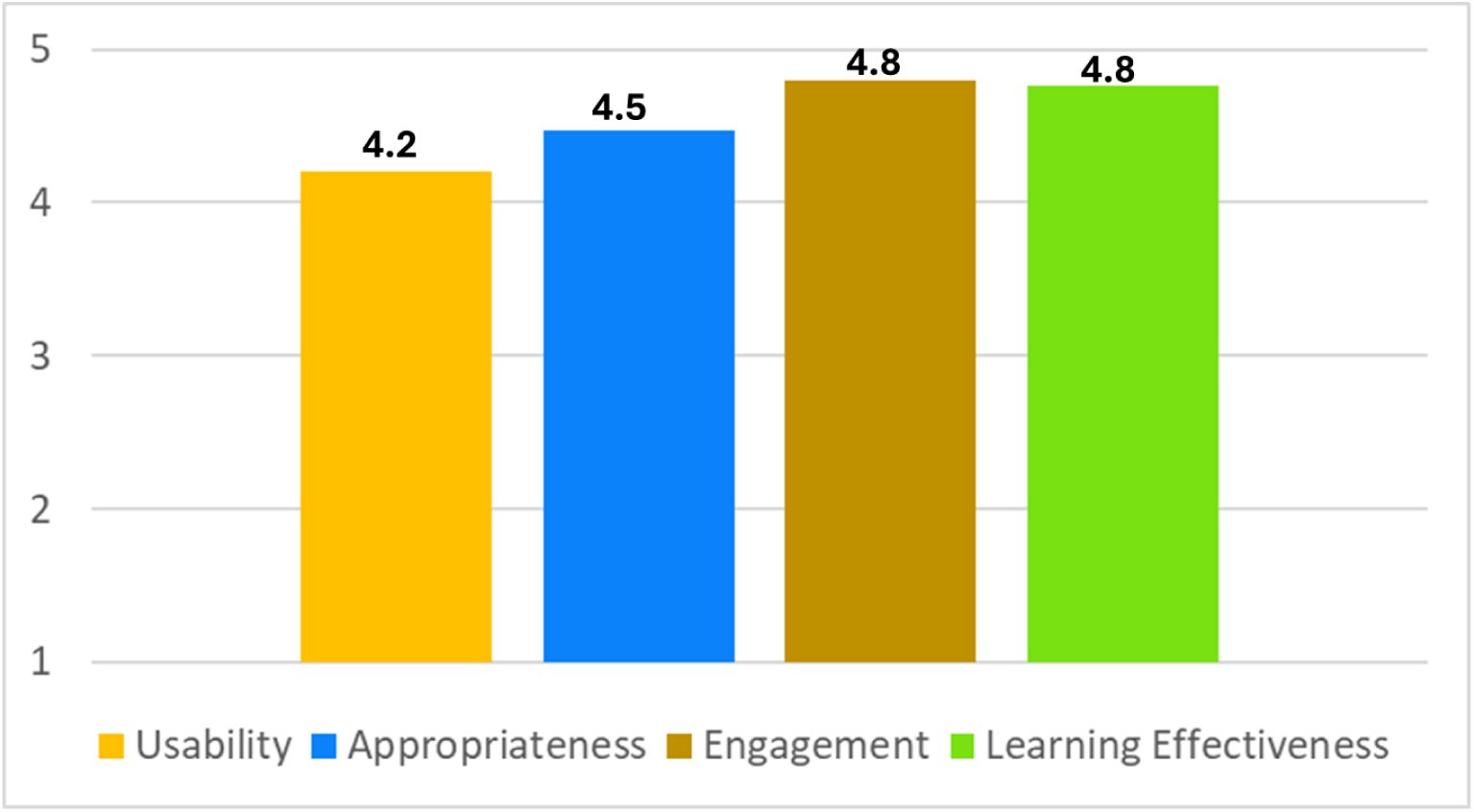
Feedback from students’ questionnaire.

## Discussion

Virtual reality has emerged as a transformative technology in education and training in universities and medical schools in recent years (Olasky et al., 2015; Pottle, 2019). In traditional university teaching, students primarily rely on classroom lectures, video presentations, and written materials, requiring them to mentally reconstruct abstract mechanisms and dynamic processes. This approach can be particularly challenging for several subjects, including physics, anatomy and physiology. In this study, we designed and developed SIGHT, an immersive VR educational platform, implemented for use with Meta Quest 3. SIGHT is based on virtual, first-person experience of healthy and defected perception, to facilitate and enhance learning of complex phenomena and cultivate empathy with the patient. The first two complementary modules of SIGHT regard the physics and functional neuroanatomy of human vision, addressing both its optical principles and the underlying neural pathways. Through interactive simulation, the optics module enables students to actively explore lens types, calculate dioptric power, understand and correct refractive defects such as myopia, presbyopia and astigmatism. The functional neuroanatomy module allows learners to navigate visual pathways in 3D and experience simulated lesions to better comprehend structure-function relationships, followed by an embedded assessment to reinforce learning outcomes. Preliminary student feedback informed further refinements of the application, improving usability and pedagogical clarity.

The goal of the present work was to move beyond passive knowledge acquisition by enabling students to directly experience these complex concepts through immersive VR. By interacting with 3D models and simulated physiological and physical processes, learners are able to actively explore, manipulate variables, and engage in problem solving within a controlled immersive environment (Dimiter Velev, 2017). Indeed, simulation models have been previously used to teach complex physiology-related concepts such as cerebral blood flow (Giannessi et al., 2008). Moreover, it has been noted that VR immersive learning experiences promote students’ cognitive engagement and exert a substantial motivational influence, particularly relevant when learners are required to engage with complex and abstract materials (Papanastasiou et al., 2019). Motivation plays a fundamental role in sustaining attention, promoting persistence in the face of cognitive difficulty, and encouraging deeper processing strategies that support durable learning. Immersive technologies can enhance learners’ sense of presence, agency, and active involvement, thereby fostering intrinsic motivation and curiosity-driven exploration.

There is considerable evidence that active learning strategies can improve teaching of concepts of physics, including optics (Sokoloff, 2016). Indeed, VR and augmented reality approaches are reshaping the traditional teaching and learning paradigms (Crogman et al., 2025), in particular in physics (Bogusevschi et al., 2020; Grivokostopoulou et al., 2017). Therefore, we designed the optics module within SIGHT to provide 3D visualization and active manipulation of lenses, often impaired to those students who don’t have access to traditional experimental labs, thus facilitating the understanding of fundamental optical concepts. Furthermore, the module enables learners to experience common refractive defects and correct them by calculating and selecting lenses with appropriate dioptric power. Through this experiential approach, students can actively apply theoretical knowledge to practical scenarios, reinforcing conceptual understanding (Crogman et al., 2025; Fidan & Tuncel, 2019).

Anatomy and physiology are complex, foundational sciences and medical students are often pushed to conceptualize the complex 3D anatomy, spatial relationships and physiology from the 2D images in the traditional text books (King, 2026; Stepan et al., 2017). Therefore, there has been an increased interest in using VR strategies and softwares in this field, in particular for their 3D capabilities which can enhance student learning (King, 2026; Vue et al., 2024; Wood et al., 2025). However, while the application to anatomy is becoming quite common, this is not yet the case for physiology (King, 2026; Vertemati et al., 2024), in particular for neurophysiology. Therefore, in the functional neuroanatomy module we have reproduced the subcortical visual pathways that transmit retinal inputs to the primary visual cortex via specific sub-bundles which are spatially connected in a complex way. The students can visualize such pathways in 3D and click on different parts to learn about their role. Furthermore, students can also directly see how lesions at different levels can lead to specific patterns of visual impairment.

Indeed, an innovative aspect of SIGHT is that it provides the possibility to experience how a person with a vision defect or lesion in the visual pathway will perceive the surroundings. Furthermore, SIGHT incorporates a problem based learning approach in which students are actively engaged in challenging situations, sorting lenses, performing calculations or correcting defects, with the evaluation part being equally challenging. Such an approach has been found to have a positive impact on the students’ learning (Aslan & Duruhan, 2021).

Overall, the preliminary student feedback pointed to a very positive experience with SIGHT. Most of the students agreed that SIGHT was easy to use, it helped them better understand complex concepts and it made learning more engaging than a traditional classroom lecture. Other VR related studies in medical education have reported positive feedback regarding VR (Zhao et al., 2021). At present, we are introducing SIGHT modules into our lectures, within the International Medical School and Medical Biotechnology and Molecular Medicine courses at the University of Milano. This will also give the opportunity to quantitatively assess learning outcomes and compare SIGHT effectiveness with traditional instructional methods. Indeed, because of the small sample size of the pilot student group, a larger and more diverse group will be needed. Moreover, we are currently designing new modules on other biomedical topics where the first person approach of SIGHT represents an added value for educational purposes.

## Conclusion

In this study, we developed SIGHT, an immersive virtual reality learning tool, combining experiential learning and direct testing of healthy and defected human perceptions. Two modules, integrating optics and functional neuroanatomy of vision, have been designed, developed and tested so far. The physics module enables students to use different lens types, estimate their dioptric power and identify and correct refractive errors such as myopia, presbyopia, and astigmatism. The neurophysiology module provides an interactive exploration of the visual pathways, allowing learners to visualize its components, simulate lesions, and observe their effects on visual perception. Although SIGHT was originally conceived for medical education, it is broadly applicable to students across health professions degree programs, as well as life science disciplines.

## Acknowledgments

We acknowledge useful discussions and feedback from Mission Brain students and several colleagues, including Massimo Locati, Laura Morelli, Cristina Tringali, Anna Marozzi, Luca Rossetti, Edoardo Villani, Paolo Ferri, Emanuele Mocciardini, Gabriella Cerri, Claudia Dellavia, Guglielmo Puglisi, Maurizio Vertemati, Anna Pistocchi, Massimo Aureli, Alex Pezzotta, Antonella Lotti and Ivano Eberini. Mirko Bove, Gisella Rossini, Maria Vittoria Ausilio and Alessandro Muiesan are also thanked for their precious technical support.

## Author’s contributions

Zulkifal Malik: Conceptualization, Visualization, Figures preparation, Writing – original draft, Luca Fornia: Conceptualization, Writing – review & editing, Josef Grunig: Conceptualization, Software, Writing – review & editing Davide Scalisi: Conceptualization, Software, Writing – review & editing, Federica Marchesi: Conceptualization, Funding acquisition, Writing – review & editing, Giuliano Zanchetta: Conceptualization, Supervision, Funding acquisition, Writing – review & editing.

## Declaration of using AI-assisted technologies

During the preparation of this work, the authors used ChatGPT (OpenAI) and Perplexity to improve the clarity and readability of some parts of the manuscript. After using these tools, the authors carefully reviewed and edited all content and they take full responsibility for the content of the publication.

## Data availability

The data that support the findings of this study are available from the corresponding author, G.Z., upon reasonable request.

## Ethical approval

All data used in this project were collected according to the rules and GDPR regulations provided by University of Milano. Informed consent was obtained from all students who participated in the questionnaire. This activity did not involve clinical intervention, collection of sensitive personal data, or any procedures that could pose risk to participants. All procedures were conducted in accordance with the principles of the Declaration of Helsinki. Data were used for educational purposes and approval from the local ethics committee of the University of Milano was therefore not required.

## Supplementary data for

**Suppl. Figure 1.**
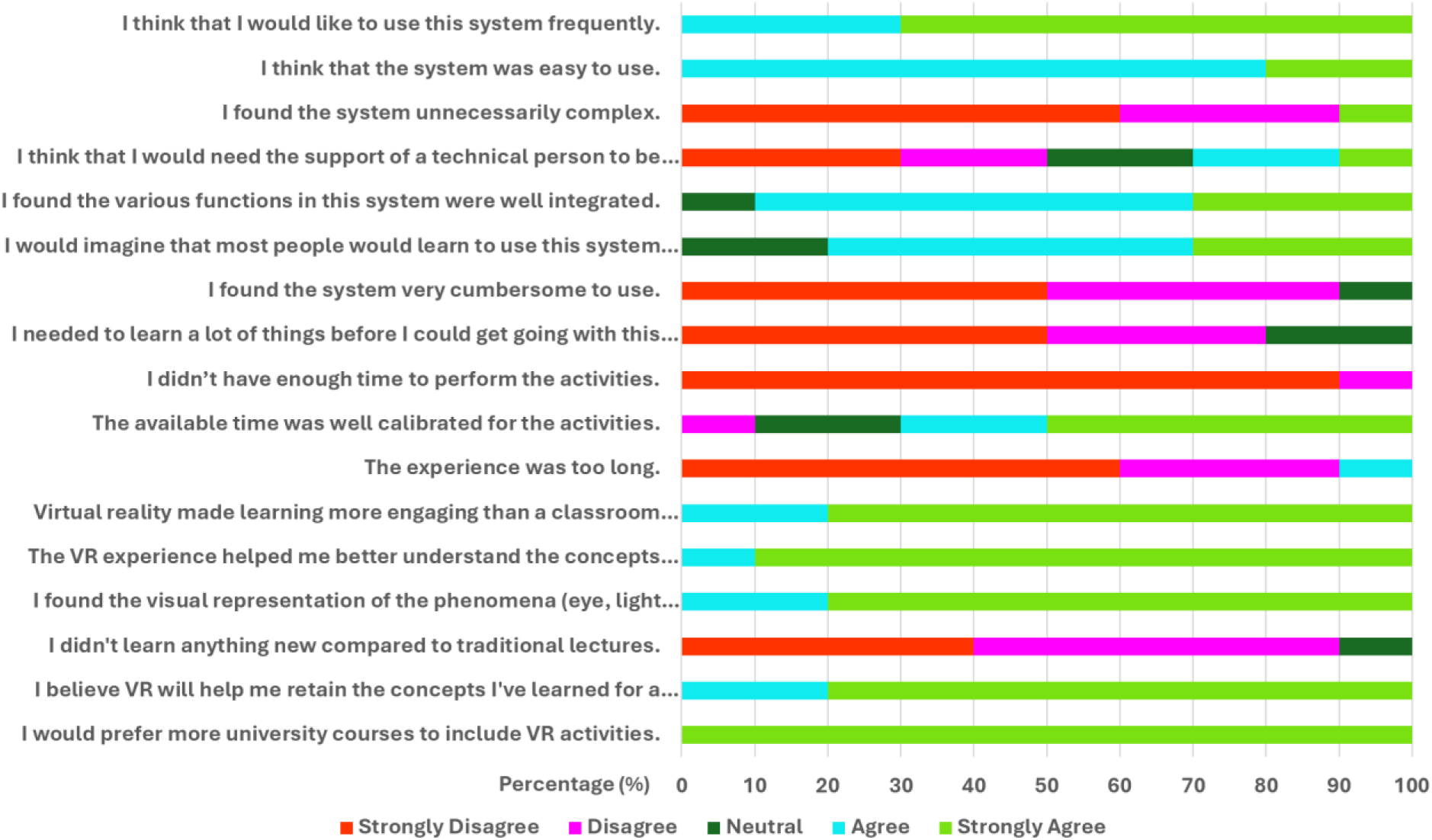
Students responses to feedback questionnaire (individual questions).

**Suppl. Table 1.** Hotspots of the visual pathway in functional neuroanatomy of vision module and their corresponding information.

| Hotspot No. | Name | Description |
| --- | --- | --- |
| 1 | Retina (Right and Left Eye) | The retina of each eye is divided into four quadrants—upper and lower, nasal and temporal—by imaginary horizontal and vertical lines passing through the center of the fovea. Retinal ganglion cells gather visual signals from photoreceptors in all these quadrants before sending them towards the brain. |
| 2 | Optic Nerve (Right and Left) | The retina of each eye is divided into four quadrants—upper and lower, nasal and temporal—by imaginary horizontal and vertical lines passing through the center of the fovea. Retinal ganglion cells gather visual signals from photoreceptors in all |
|  |  | these quadrants before sending them towards the brain |
| 3 | Optic Chiasm | The axon fibers in the right and left optic nerves reach optic chiasm. Here, fibers from the nasal half of each retina cross to the opposite side, while temporal fibers stay on the same side. This crossing allows both brain hemispheres to receive and share visual information from both eyes. |
| 4 | Optic Tract (Right and Left) | From the optic chiasm, axon fibers continue toward the thalamus as the optic tracts in both hemispheres. Each tract contains fibers from both the nasal and temporal quadrants of the retina and carries visual information from both eyes |
| 5 | Lateral Geniculate Nucleus (Right and Left) | The axons in each optic tract mainly project to the lateral geniculate nucleus of the thalamus. Fibers from the right optic tract connect to the right LGN, and fibers from the left tract connect to the left LGN, where visual information is organized before reaching the cortex |
| 6 | Optic Radiations (Right and Left) | In both hemispheres, the axonal fibers leave the lateral geniculate nucleus as optic radiations. These fibers carry visual signals to the visual cortex. |
| 7 | Meyer's Loop or Inferior Optic Radiations (Right and Left) | Some fibers of the optic radiations form a curve called Meyer's loop, passing through the temporal lobe before reaching the visual cortex in both hemispheres. These fibers carry visual information from the upper part of the visual field of both eyes |
| 8 | Dorsal Bundle or Superior Optic Radiations | Some optic radiation fibers pass beneath the parietal lobe on their way to the visual cortex in both hemispheres. These fibers carry visual information from the lower part of the visual field of both eyes |
| 9 | Visual Cortex (V1) Above the Calcarine Sulcus (Cuneus Region) | In both hemispheres, fibers from the inferior visual field project to the visual cortex above the calcarine sulcus, in the cuneus region. This area processes input from the lower visual field of both eyes |
| 10 | Visual Cortex (V1) Below the Calcarine Sulcus (Lingual Gyrus) | In both hemispheres, fibers from the superior visual field project to the visual cortex below the calcarine sulcus, in the lingual gyrus. This area processes input from the upper visual field of both eyes |

**Suppl. Table 2.** Lesions of the visual pathway and their corresponding defects. Lesions can be induced in the left or right side of the visual pathway.

| Lesions | Defect Visual Field | Description |
| --- | --- | --- |
| 1 | Central Scotoma (Left and Right) | A central scotoma is a blind spot in the center of the visual field. It usually results from damage to the macula or the optic nerve, affecting the ability to see fine details directly in front of you. |
| 2 | Monocular Vision Loss (Left and Right) | Damage to one of the optic nerves before it reaches the optic chiasm results in loss of vision that is limited to the eye of origin. |
| 3 | Bitemporal Hemianopia | Bitemporal hemianopia happens when the center of the optic chiasm is damaged, often by a pituitary tumor. This affects the crossing nasal fibers from both eyes, leading to loss of vision in the outer, or temporal, visual fields, because the nasal retina receives light from the outer(temporal) part of the visual field. |
| 4 | Contralateral Homonymous Hemianopia (Left and Right) | Damage to the visual pathway on one side, whether in the optic tract, optic radiations, or visual cortex, causes loss of the same half(contralateral) of the visual field in both eyes. For example, damage on the right side leads to loss of vision in the left half of the visual field in both eyes. |
| 5 | Contralateral Superior Quadrantopia (Left and Right) | When Meyer's loop in the temporal lobe is damaged, vision is lost in the upper quarter of the visual field on the opposite side. For example, damage to the right temporal lobe causes loss of the upper left visual field in both eyes. |
| 6 | Contralateral Inferior Quadrantopia (Left and Right) | When the superior optic radiations in the parietal lobe are damaged, vision is lost in the lower quarter of the visual field on the opposite side. For example, damage to the right parietal lobe causes loss of the lower left visual field in both eyes. |
| 7 | Contralateral Homonymous Hemianopia with Macular Sparing (Left and Right) | When the primary visual cortex is damaged on one side, vision is lost in the opposite half of the visual field. The central (macular) vision may be preserved if the occipital pole remains intact. For example, right occipital cortex damage causes left visual field loss, but central vision can still be seen |

